# Using nucleocapsid proteins to investigate the relationship between SARS-CoV-2 and closely related bat and pangolin coronaviruses

**DOI:** 10.1101/2020.06.25.172312

**Authors:** Noah Schuster

**Affiliations:** Department of Biology, DePauw University, Greencastle, Indiana

## Abstract

An initial outbreak of coronavirus disease 2019 (COVID-19) in China has resulted in a massive global pandemic causing well over 16,500,000 cases and 650,000 deaths worldwide. The virus responsible, SARS-CoV-2, has been found to possess a very close association with Bat-CoV RaTG13 and Pangolin-CoV MP789. The nucleocapsid protein can serve as a decent model for determining phylogenetic, evolutionary, and structural relationships between coronaviruses. Therefore, this study uses the nucleocapsid gene and protein to further investigate the relationship between SARS-CoV-2 and closely related bat and pangolin coronaviruses. Sequence and phylogenetic analyses have revealed the nucleocapsid gene and protein in SARS-CoV-2 are both closely related to those found in Bat-CoV RaTG13 and Pangolin-CoV MP789. Evidence of recombination was detected within the N gene, along with the presence of a double amino acid insertion found in the N-terminal region. Homology modeling for the N-Terminal Domain revealed similar structures but distinct electrostatic surfaces and topological variations in the β-hairpin that likely reflect specific adaptive functions. In respect to SARS-CoV-2, two amino acids (S37 and A267) were found to exist only in its N protein, along with an extended β-hairpin that bends towards the nucleotide binding site. Collectively, this study strengthens the relationship among SARS-CoV-2, Bat-CoV RaTG13, and Pangolin-CoV MP789, providing additional insights into the structure and adaptive nature of the nucleocapsid protein found in these coronaviruses. Furthermore, these data will enhance our understanding of the complete history behind SARS-CoV-2 and help assist in antiviral and vaccine development.

## Introduction

In December 2019, an outbreak of an unknown disease occurred in Wuhan, China.^1^ Soon after, this illness was named coronavirus disease 2019 (COVID-19) and defined as a respiratory infection caused by severe acute respiratory syndrome coronavirus 2 (SARS-CoV-2), with patients displaying symptoms that manifested as mild or severe cases of pneumonia.^1,2^ Initial cases showed very close associations with the local animal market in Wuhan, which contained both live and dead animals, suggesting a spillover event occurred from an unidentified animal source into the human population.^2^ Following its emergence in China, SARS-CoV-2 has managed to spread across six continents and result in over 16,500,000 confirmed cases and 650,000 deaths worldwide.^3^ As of now, there is still no available vaccine to combat the disease, and while the number of cases and deaths continue to grow, there is a huge need to continue research that studies every aspect of the virus, including its phylogeny and evolution.

Coronaviruses (CoVs) are diverse single-stranded positive-sense RNA viruses with genomes ~30 kilobases in length and belong to the subfamily Coronavirinae in the family Coronaviridae of the order Nidovirales. Furthermore, there are four recognized and distinct genera in Coronavirinae: *Alphacoronavirus*, *Betacoronavirus*, *Deltacoronavirus*, and *Gammacoronavirus*.^4^ Additionally, these four genera then split into a variety of distinct subgenera (https://talk.ictvonline.org/ictv-reports/). Officially, SARS-CoV-2 has been ranked within the genus *Betacoronavirus* and the subgenus *Sarbecovirus*, which are the same groups that contain severe acute respiratory syndrome coronavirus (SARS-CoV).^5^ An overall 96% genome identity revealed the closest relative to SARS-CoV-2 was Bat-CoV RaTG13, followed by an 88% genome identity with bat-SL-CoVC45 and bat-SL-CoVZXC21.^6^ In relation to SARS-CoV, only 79% genome identity was found to indicate a more genetically distant relationship exists between each virus.^6,7^ Recently, CoVs that were isolated from pangolins were found to have a 90-91% genome identity with SARS-CoV-2.^8,9^ In the spike protein, the receptor-binding domain (RBD) in SARS-CoV-2 held greater similarity with pangolin CoVs in six key RBD residues unlike in Bat-CoV RaTG13 which only shares a single residue.^10^ Therefore, it has been hypothesized that recombination occurred during the evolution of pangolin CoVs and Bat-CoV RaTG13, contributing to the development of SARS-CoV-2.^8,9,10^ Additionally, these particular CoVs tend to form a separate clade in the *Sarbecovirus* subgenus when viewed on either genome-based or gene-based phylogenetic trees, furthering the likelihood of a shared ancestry that is more distinct from other distantly related CoVs.^8,11,12^

Although phylogeny for SARS-CoV-2 is largely established, the complete evolutionary history for the virus is still in question. A recent investigation that focused on the molecular evolution of SARS-CoV-2 indicated majority of the open-reading frames undergo purifying selection, and there exists little evidence of positive selection.^13^ Furthermore, bats are natural reservoirs that possess a variety of CoVs.^14,15^ With greatest similarity to Bat-CoV RaTG13 found in horseshoe bats (*Rhinolophus affinis*), SARS-CoV-2 likely has its origin in bats as well. While bats serve as the natural reservoir, the identity of an intermediate host is largely unresolved. A range of conflicting studies have come out to suggest, but not definitely prove, that snakes, turtles, yaks, and/or Malayan pangolins (*Manis javanica*) could potentially be the intermediate host(s) for the virus.^8,9,16–18^ Even if one of these animals is the intermediate host, SARS-CoV-2 has yet to be isolated from any of them.^19^ Nonetheless, many CoVs go through an intermediate host prior to reaching their intended host; therefore, it seems unlikely that SARS-CoV-2 would fail to possess an intermediate host of some kind.^15,19^

In the remaining one-third of all CoV genomes, there are several open-reading frames near the 3’-terminus that encode four main structural proteins: the nucleocapsid (N), membrane (M), envelope (E), and spike (S) proteins.^4,20^ The N protein is multifunctional and involved in a variety of processes such as genome encapsidation and circularization, transcription, RNA packaging, genome transport to viral budding sites, repressing interferon production, cell cycle inhibition, and blocking RNA interference.^20–23^ Furthermore, the N protein is also highly immunogenic and largely overexpressed during replication, thereby capable of inducing large-scale immune responses against CoVs.^24,25^ Overall, the N protein possesses two highly conserved domains: the N-Terminal Domain (NTD) and C-Terminal Domain (CTD).^23,26^ Between each domain is an intrinsically disordered but flexible region known as the Linker Region (LKR) that is rich in serine and arginine residues.^26^ There are two additional intrinsically disordered regions located at each terminus: the N-Terminal Region (NTR) and C-Terminal Region (CTR).^26^ Each disordered region can interact with either the genome or viral proteins following the phosphorylation of residues that induce conformational changes to promote high affinity binding.^26,27^ In particular, the NTD is divergent in sequence and length; however, it has a β-sheet core along with a highly electropositive groove and β-hairpin extension that binds RNA to neutralize negative charges found within the phosphate-sugar backbone.^26^ For majority of CoVs their NTDs are structurally similar; nevertheless, multiple electrostatic surface comparisons have revealed uniquely arranged charge distributions, suggesting that CoV NTDs utilize distinct modes of RNA-binding.^26,28^ In terms of function, the NTD serves as scaffolding to assist in genome packaging and virion assembly.^28,29^

From an evolutionary standpoint, the N gene in SARS-CoV-2 was found to be under strong selective pressure and displayed higher genetic diversity, increased host adaptation, and was prone to natural selection.^30^ The structure of the dimerization domain in the SARS-CoV N protein has also helped to provide a link between Coronaviridae and Arteriviridae, highlighting the importance of structural data in assessing the relationship between different viruses.^31^ Since the N protein is largely conserved and found to be evolving faster than other CoV proteins, it can serve as a model for establishing phylogenetic relationships, assessing mutations, calculating divergence times, and determining if recombination events occurred among specific CoVs.^14,30^

Currently, there is a limited amount of research focusing exclusively on the N protein in SARS-CoV-2. Herein, this study relies on a strictly bioinformatic and homology modeling approach in which both N gene and protein sequences are used to further investigate the characteristics shared between SARS-CoV-2 and a set of closely related bat and pangolin CoVs.

## Materials and Methods

### 1. Sequence Acquisition and Identity Analysis

Nucleocapsid gene and protein sequences in SARS-CoV-2, pangolin CoVs, and bat CoVs were retrieved from the NCBI RefSeq, Nucleotide, and Protein databases. Percent identity was calculated by MatGAT v2.01 to determine the shared identity for each sequence type.^32^ For nucleotide identity, the first gap and extending gap penalties were set at 16 and 4, respectively. As for amino acid identity, a PAM250 scoring matrix was used along with the first gap and extending gap penalties set at 12 and 2, respectively.

### 2. Phylogenetic and Recombination Analysis

Sequences were aligned using MUSCLE on the MEGA-X v10.1.7 software.^33,34^ The default alignment settings were retained, except clustering methods which were both changed to neighbor joining. In addition, nucleotide sequences were aligned by codons. An individual gene tree and protein tree were constructed using different maximum likelihood methods and visualized on MEGA-X.^35^ While constructing both trees, uniform rates and partial deletion were used. The heuristic model was nearest-neighbored-interchange and the initial tree was automatically made with the default option NJ/BioNJ. These trees were not rooted with an outgroup.

Based on the protein tree, one sequence was chosen from each split. Using those sequences, nucleotide and amino acid identities for the NTD and CTD were calculated by MatGAT v2.01 to evaluate the identity shared among each domain. Individual gene and protein trees were also constructed for each domain in the same manner as previously described. Gene sequences were aligned using MUSCLE, after which the alignment was used by SimPlot v3.5.1 to generate a nucleotide similarity plot in detecting for potential recombination events.^36^ The similarity plot was computed with a Kimura (2-parameter) distance model and similarity scores were calculated using a 250 bp sliding window and step of 10 bp, along with a 50% consensus threshold.

### 3. Disorder and Phosphorylation Site Predictions

Calculated predictions for protein disorder were determined using PrDOS, IUPred2, ANCHOR2, DISOPRED3, and PONDR.^37–40^ PrDOS was run using the standard 5.0% false positive rate. IUPred2 and ANCHOR2 were both run using the long sequence option with default settings. DISOPRED3 was run using default settings. PONDR was run using default settings and the VLXT results were obtained. Predictions for disorder were normalized on a 0.0-1.0 scale for each program, with values greater than 0.5 to indicate residues with a tendency to be considered disordered. This prediction method is similar to one previously described; therefore, it is important to note this method provides a conservative estimate of disorder and prevents the possibility of over-estimating the disorder for any specific residue.^41^

To determine which residues are most likely to be phosphorylated, the DEEP online server was used (http://www.pondr.com/cgi-bin/depp.cgi). Phosphorylation predictions were normalized on a 0.0-1.0 scale, with values greater than 0.5 to indicate residues having the potential of being phosphorylated. This method does not guarantee phosphorylation occurs but provides scoring for the likelihood that phosphorylation would occur for any given serine, threonine, and/or tyrosine residue within a protein.

### 4. Homology Modeling

Homology models were built using the NTD in SARS-CoV (PDB: 2OFZ) and carried out using the MODELLER v9.23 software.^42,43^ Ten models were built for each sequence and the model presenting with the lowest discrete optimized protein energy (DOPE) was chosen. Models were then evaluated using Verify-3D, ERRAT, and PROCHECK on the SAVES v5.0 online sever (https://servicesn.mbi.ucla.edu/SAVES/). Verify-3D measures the compatibility between a model and its own amino acid sequence, calculating scores ≥ 80% to indicate ideal compatibility.^44,45^ ERRAT measures the quality of non-bonded interactions between atoms, producing scores ≥ 90 to indicate ideal quality.^46^ PROCHECK generates a Ramachandran plot to verify torsion angles and determines if residues are properly built; furthermore, at least ≥ 90% of these residues should fall within the favorable region.^47,48^

To ensure a model had the most stable conformation, regions that presented with either low or moderate scoring were subjected to energy minimization using UCSF Chimera.^49^ First, hydrogen atoms were added. Then, charges were assigned using the parameters designated for standard residues in the AMBER ff14SB option. Lastly, the steepest descent and conjugate gradient technique was employed with 100 and 10 steps, respectively. The stereochemical quality of the model was determined again with Verify-3D, ERRAT, and PROCHECK. The model was reliable if it presented with improved values than previously determined by each program.

### 5. Comparing Models

Once NTD models were constructed, they were superimposed on UCSF Chimera to determine root-mean-square deviation (RMSD) values.^50^ Parameters used for superimposition were defined by the Needleman-Wunsch algorithm and a BLOSUM62 scoring matrix, along with a residue-residue distance cut-off of 5.0 Å. The gap open and extension penalties were 12 and 1, respectively. All other default settings were maintained. If the RMSD between two models was low, they were viewed as structurally similar. The structural distance measure (SDM) and Q-Score were used to evaluate the entire structural similarity among all the NTD models.^51,52^ Topological variations within secondary structures, motifs, loops, and/or coils were considered as well. Lastly, electrostatic surface maps were determined by UCSF Chimera and visualized according to the default columbic surface coloring schemes.

## Results

A total of 10 nucleocapsid gene and protein sequences were used in this study. Among these sequences, six were from pangolin CoVs, three were from bat CoVs, and the remaining one was from SARS-CoV-2 (Table S1). To ensure clarity and specificity, the isolate name was used to differentiate between the multiple pangolin and bat CoVs in the sequence dataset. Notably, five of the six pangolin CoVs were isolated between 2017-2018.

The SARS-CoV-2 N gene shares a high sequence identity with Bat-CoV RaTG13 (97%) and Pangolin-CoV MP789 (96%); furthermore, N genes found in pangolin CoVs isolated between 2017-2018, along with bat-SL-CoVZXC21 and bat-SL-CoVZC45, were found to share lower identities at 90-91% (Table S2). Similarly, the SARS-CoV-2 N protein shares a high sequence identity with Bat-CoV RaTG13 (99%) and Pangolin-CoV MP789 (97%). Moreover, the N proteins in pangolin CoVs isolated between 2017-2018, along with bat-SL-CoVZXC21 and bat-SL-CoVZC45, were also found to share lower identities at 93-94% (Table S2).

In particular, the alignment revealed that within the N-terminal region of the N proteins in SARS-CoV-2, Pangolin-CoV MP789, and Bat-CoV RaTG13, each presented with a double amino acid insertion consisting of arginine and asparagine (Figure 1; Figure S1). This insertion was observed in bat-SL-CoVZXC21 and bat-SL-CoVZC45; however, bat-SL-CoVZXC21 had two serine residues while bat-SL-CoVZC45 had arginine and serine. In addition, residues shared between pangolin and bat CoVs (P37 and Q267) were mutated to S37 and A267 in SARS-CoV-2. Six amino acids (G25, S26, P80, N345, Q349, and T379) were shared only among N proteins found in SARS-CoV-2, Bat-CoV RaTG13, and Pangolin-CoV MP789; meanwhile, two additional residues (K65 and D128) were also found but shared only between Bat-CoV RaTG13 and SARS-CoV-2. For pangolin CoVs isolated between 2017–2018, many conserved amino acid groups were detected throughout the alignment (Figure S1). Lastly, no sites were found shared only between Pangolin-CoV MP789 and SARS-CoV-2.

**Figure 1.**
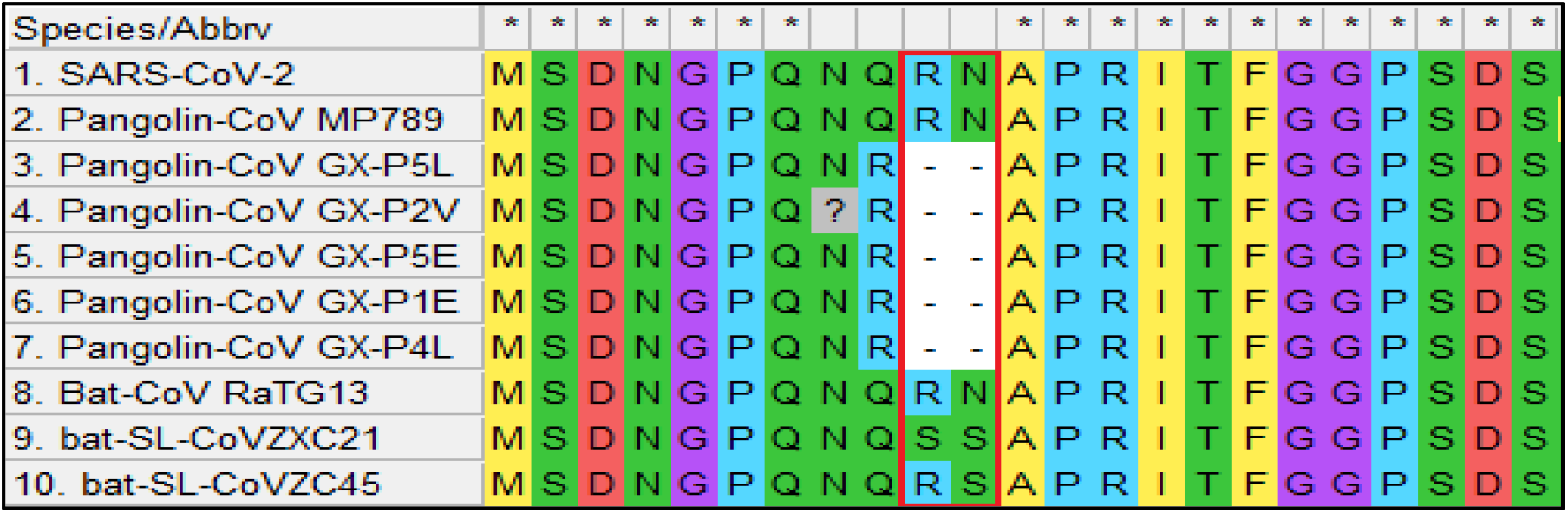
Screen capture of the MUSCLE alignment for the start of the N protein. The red box indicates the insertion region present in SARS-CoV-2, Pangolin-CoV MP789, Bat-CoV RaTG13, bat-SL-CoVZXC21, and bat-SL-CoVZC45.

Phylogenetic analysis suggests the N gene in SARS-CoV-2 is genetically closer to the N gene found in Bat-CoV RaTG13, followed by a more distant relationship to the N gene found in Pangolin-CoV MP789 (Figure 2A). In addition, the N genes found in pangolin CoVs that were isolated between 2017-2018 formed a polytomy while bat-SL-CoVZXC21 and bat-SL-CoVZC45 functioned as sister taxa that were distant from all other N genes in the tree (Figure 2A). Phylogenetic analysis of the N protein revealed the same groupings of CoVs as presented in the gene tree; however, the topology is different (Figure 2B). The protein tree shows bat-SL-CoVZXC21 and bat-SL-CoVZC45 in closer relation to SARS-CoV-2, whereas both CoVs in the gene tree held a lesser relationship to SARS-CoV-2. Conversely, the gene tree has pangolin CoVs isolated between 2017-2018 in closer relation to SARS-CoV-2, unlike the protein tree where they have a lesser relation to SARS-CoV-2 (Figure 2).

**Figure 2.**
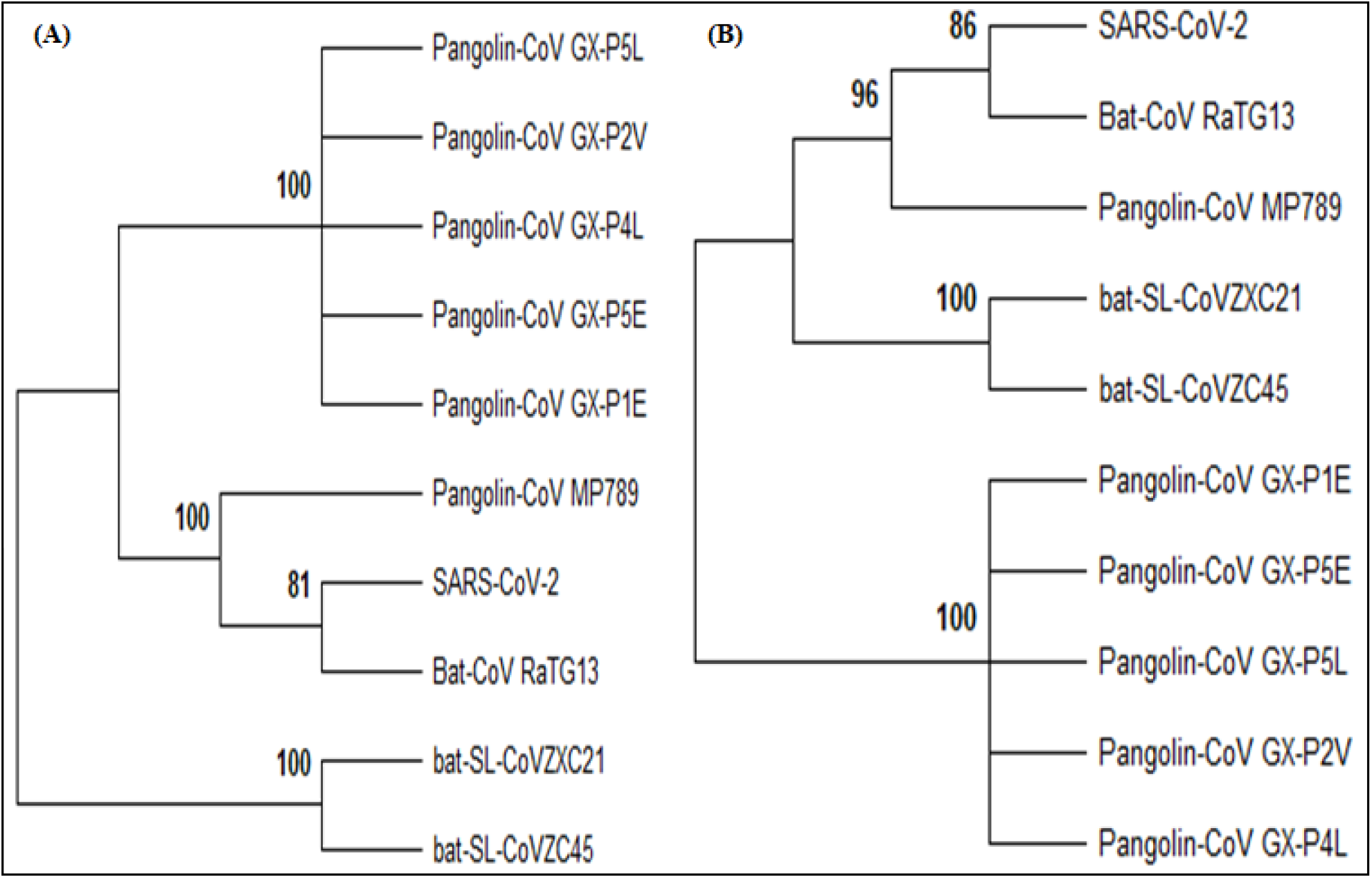
(A) Phylogenetic analysis of N gene sequences depicting evolutionary relationships observed among SARS-CoV-2, pangolin CoVs, and closely related bat CoVs. The phylogeny was inferred using the Jukes-Cantor substitution model. A log likelihood of −3,096.80 was determined with bootstrap values calculated out of 500 replicates and nodes with < 70.00% support collapsed. (B) Phylogenetic analysis of N protein sequences depicting evolutionary relationships observed among SARS-CoV-2, pangolin CoVs, and closely related bat CoVs. The phylogeny was inferred using the JTT matrix-based substitution model. A log likelihood of −1,560.88 was determined with bootstrap values calculated out of 500 replicates and nodes with < 70.00% support collapsed.

Based on the phylogenetic positions of each N protein, one representative sequence was selected from each split in the tree for additional analysis. From this, N proteins that belonged to SARS-CoV-2, Bat-CoV RaTG13, Pangolin-CoV MP789, bat-SL-CoVZXC21, and Pangolin-CoV GX-P5L were chosen. It was found that the NTD in SARS-CoV-2 is 100% identical to the NTD in Bat-CoV RaTG13 while the CTD in SARS-CoV-2 is 99% identical to the CTD in Bat-CoV RaTG13 (Table 1). The N protein domains in Pangolin-CoV MP789 shared a very high identity with SARS-CoV-2 at 98% for the NTD and CTD as well (Table 1). In fact, all of the selected N protein sequences expressed high levels of identity for the NTD and CTD with the lowest values being only 94% and 96%, respectively (Table 1). Nucleotide sequences corresponding to each domain followed similar trends as well; however, it is interesting that the NTD holds noticeably higher nucleotide identities than does the CTD (Table 1). Furthermore, phylogenetic analysis of nucleotide sequences revealed the NTD in SARS-CoV-2 is genetically similar to the NTDs found in Bat-CoV RaTG13 and Pangolin-CoV MP789, whereas the analysis of amino acid sequences revealed the NTD in SARS-CoV-2 is closer in relation to the NTD found in Bat-CoV RaTG13 (Figure S2 A,B). Conversely, phylogenetic analysis of nucleotide sequences revealed the CTD in SARS-CoV-2, Bat-CoV RaTG13, and Pangolin-CoV MP789 are genetically similar, with the same topology observed in the protein tree as well (Figure S2 C,D). In regard to both Pangolin-CoV GX-P5L and bat-SL-CoVZXC21, they either formed phylogenetically distinct sister taxa or independent lineages based on their NTDs and CTDs (Figure S2).

**Table 1.** Regions defined as the NTD and CTD within the N gene and protein. In addition, the nucleotide and amino acid identities are shown for both domains when set-up against SARS-CoV-2.

|  | NTD Regions |  | NTD Identities (%) |  | CTD Regions |  | CTD Identities (%) |  |
| --- | --- | --- | --- | --- | --- | --- | --- | --- |
|  | Gene | Protein | Nucleotide | Amino Acid | Gene | Protein | Nucleotide | Amino Acid |
| Pangolin-CoV MP789 | 142–519 | 48–173 | 97.9 | 97.6 | 739–1,092 | 247–364 | 94.6 | 98.3 |
| Pangolin-CoV GX-P5L | 136–513 | 46–171 | 93.9 | 94.4 | 733–1,086 | 245–362 | 90.7 | 96.6 |
| bat-SL-CoVZXC21 | 142–519 | 48–173 | 93.9 | 96.0 | 739–1,092 | 247–364 | 88.4 | 96.6 |
| Bat-CoV RaTG13 | 142–519 | 48–173 | 98.9 | 100.0 | 739–1,092 | 247–364 | 95.5 | 99.2 |

A similarity plot reveals that sequence identities are not homogenous among N genes found within SARS-CoV-2, Bat-CoV RaTG13, Pangolin-CoV MP789, bat-SL-CoVZXC21, and Pangolin-CoV GX-P5L. The complete span of the N gene found in Bat-CoV RaTG13 shares higher nucleotide identity with Pangolin-CoV MP789 and SARS-CoV-2 than with either bat-SL-CoVZXC21 or Pangolin-CoV GX-P5L (Figure 3). However, nucleotides 485-565 found in Pangolin-CoV MP789, bat-SL-CoVZXC21, and Bat-CoV RaTG13 share nearly the same identity to each other and with SARS-CoV-2 (Figure 3). Interestingly, that same region does not possess any shared identity alongside Pangolin-CoV GX-P5L. As with percent identities, this similarity plot also reveals how the NTD and CTD in Bat-CoV RaTG13 is most similar to the domains in SARS-CoV-2 (Figure 3).

**Figure 3.**
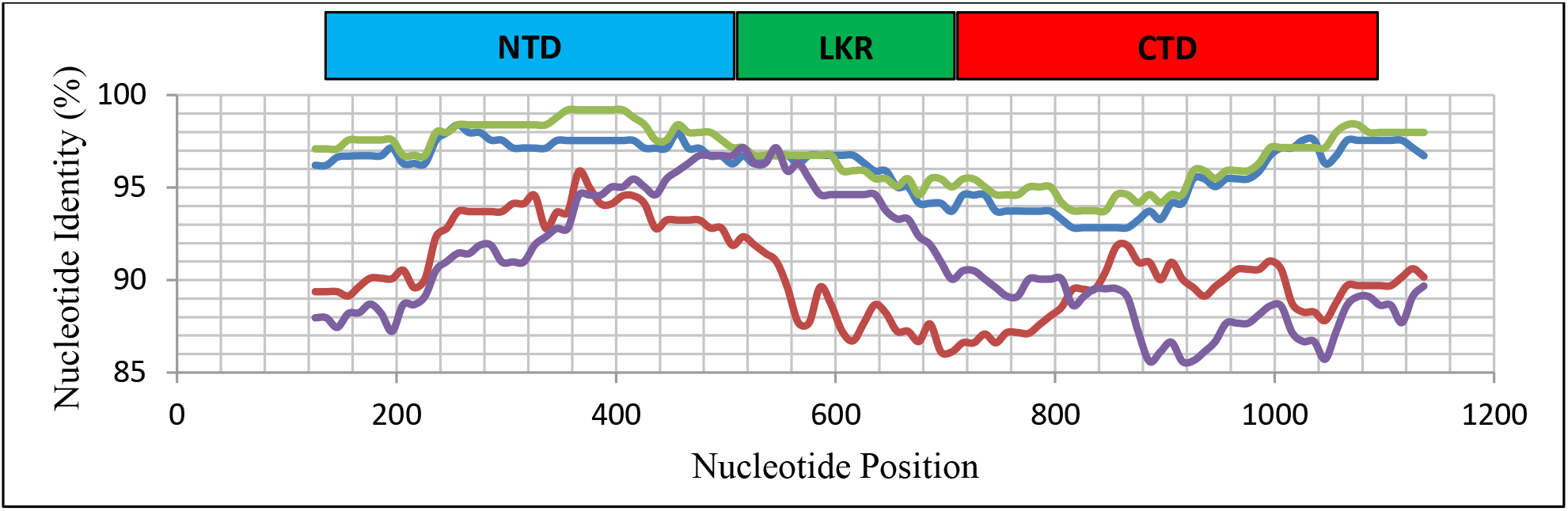
Similarity plot based on the N gene sequence from SARS-CoV-2. Subject sequences, as depicted in the plot, are from Bat-CoV RaTG13 (green), bat-SL-CoVZXC21 (purple), Pangolin-CoV MP789 (blue), and Pangolin-CoV GX-P5L (red). Nucleotide positions 1-125 and 1,160-1,260 are not shown.

Disorder predictions for the N protein revealed nearly identical values of disorder for each residue (Table S3). Higher values of disorder were observed in the NTR, LKR, and CTR; however, slight variations were observed for these regions (Figure 4). As for the NTD and CTD, both domains presented with values that were either below or near the disorder threshold; however, the beginning of the CTD showed higher values of disorder that gradually decreased when moving further into the domain (Figure 4). The detection of phosphorylation sites presented similar scoring among each N protein; however, residues with scores suggesting phosphorylation existed mainly within the NTR, LKR, and CTR (Table S4; Figure 5). Interestingly, residues found in the LKR had nearly identical scores whereas residues in the NTR and CTR presented with more varied amounts of scoring (Figure 5).

**Figure 4.**
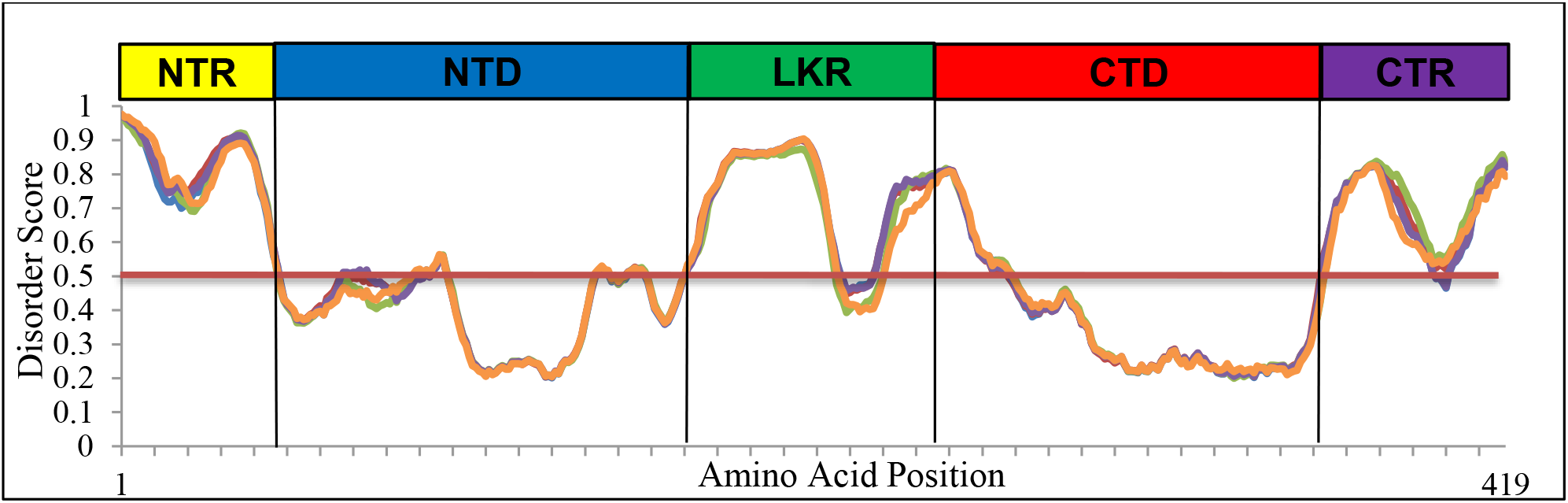
Intrinsic disorder plot for N proteins found within SARS-CoV-2 (blue), Bat-CoV RaTG13 (purple), bat-SL-CoVZXC21 (orange), Pangolin-CoV MP789 (red), and Pangolin-CoV GX-P5L (green). The red line indicates the disordered threshold and the black lines represent divisions between each different segment of the N protein. Though not indicated on the plot, the intervals on the x-axis go by units of ten.

**Figure 5.**
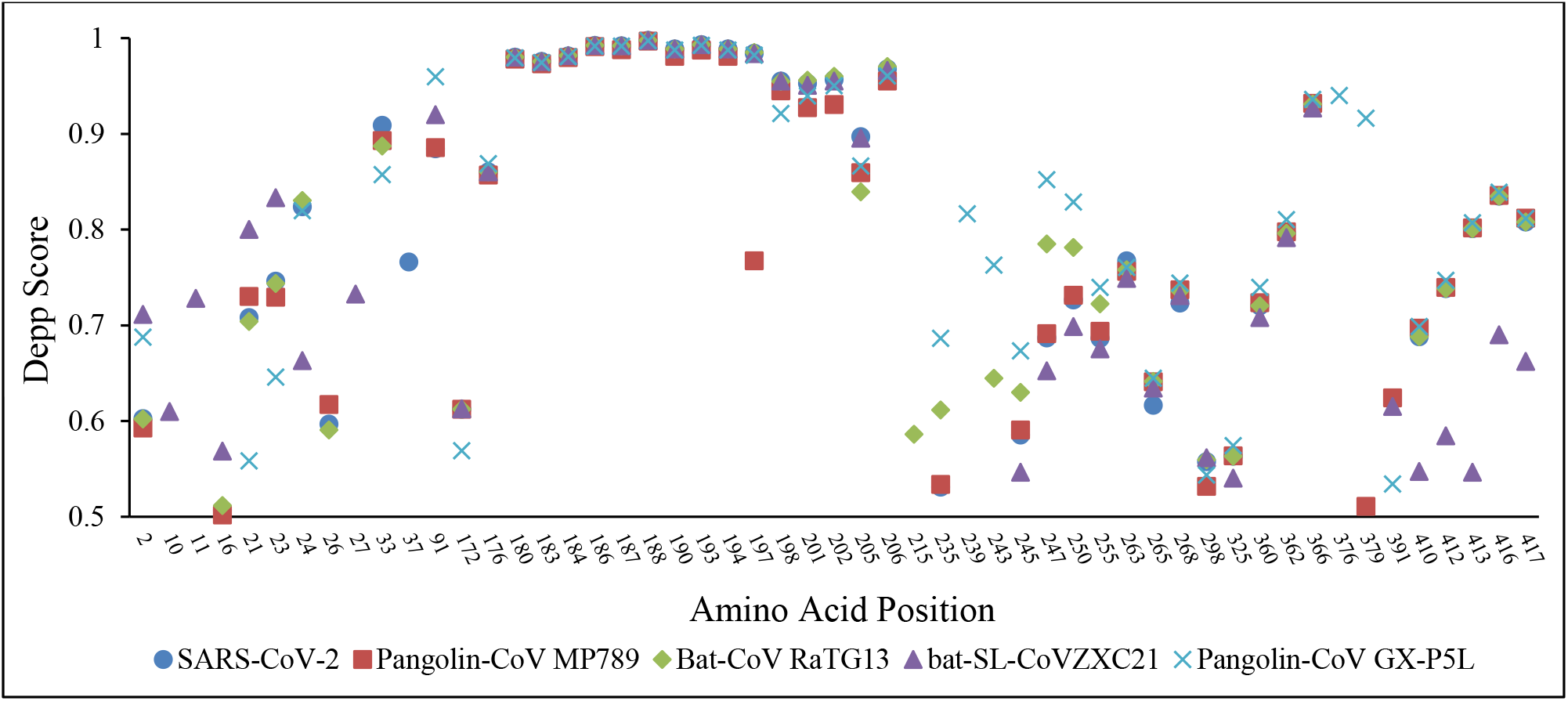
Phosphorylation plot for N proteins found within SARS-CoV-2 (blue), Bat-CoV RaTG13 (purple), bat-SL-CoVZXC21 (orange), Pangolin-CoV MP789 (red), and Pangolin-CoV GX-P5L (green). Only residues with a score greater than 0.5 are shown. The higher the score indicates a greater likelihood that a specific residue in the N protein will undergo phosphorylation.

Once homology models were constructed, they were refined and validated to meet appropriate energy and stereochemical parameters (Table S5). Models were then superimposed to yield RMSD values (Figure 4A; Table 2). From this, the NTDs in Pangolin-CoV MP789 and Bat-CoV RaTG13 were structurally similar to the NTD found in SARS-CoV-2; furthermore, the NTDs found in Pangolin-CoV GX-P5L and bat-SL-CoVZXC21 each showed greater structural similarity towards the NTD found in Pangolin-CoV MP789. The overall RMSD was 0.385, along with a calculated SDM and Q-score of 7.686 and 0.984, respectively. These values indicate that there is a significantly high structural similarity among all the constructed CoV NTDs.

**Table 2.** Matrix displaying the RMSD values shared throughout a 126 amino acid distance between NTD homology models. ‘-‘ Empty Box.

|  | 1 | 2 | 3 | 4 |
| --- | --- | --- | --- | --- |
| 1. SARS-CoV-2 | - | - | - | - |
| 2. Bat CoV RaTG13 | 0.389 | - | - | - |
| 3. bat-SL-CoVZXC21 | 0.356 | 0.504 | - | - |
| 4. Pangolin-CoV GX-P5L | 0.422 | 0.462 | 0.375 | - |
| 5. Pangolin-CoV MP789 | 0.329 | 0.429 | 0.200 | 0.289 |

In looking at specific motifs, the β-hairpin (amino acids 93–102) is where a majority of topological diversity can be observed (Figure 6B). The variations in this motif were largely present in the loop connecting both β-sheets; however, the β-sheets found in Bat-CoV RaTG13 present with a slightly different topology in comparison with the other CoV NTDs. Visually, the β-hairpin region in SARS-CoV-2 extends furthest from the protein core while also bending closest towards the nucleotide binding site. Oppositely, the β-hairpin region in Pangolin-CoV GX-P5L extends the shortest distance from the protein core. Additionally, the β-hairpin region in Bat-CoV RaTG13 appears as an intermediate that does not extend as far or as short when compared with those in SARS-CoV-2 and Pangolin-CoV GX-P5L. As for β-hairpin regions in Pangolin-CoV MP789 and bat-SL-CoVZXC21, they expressed the greatest similarity. Interestingly, the β-hairpin motif ultimately varies topologically despite homogeneity in amino identity, except at residue 94 in Pangolin-CoV GX-P5L which is valine instead of isoleucine (Figure S1). As a result of these differences, the surface features of the β-hairpin are varied, thereby presenting with differences in the distribution of positive charge as well (Figure S3).

**Figure 6.**
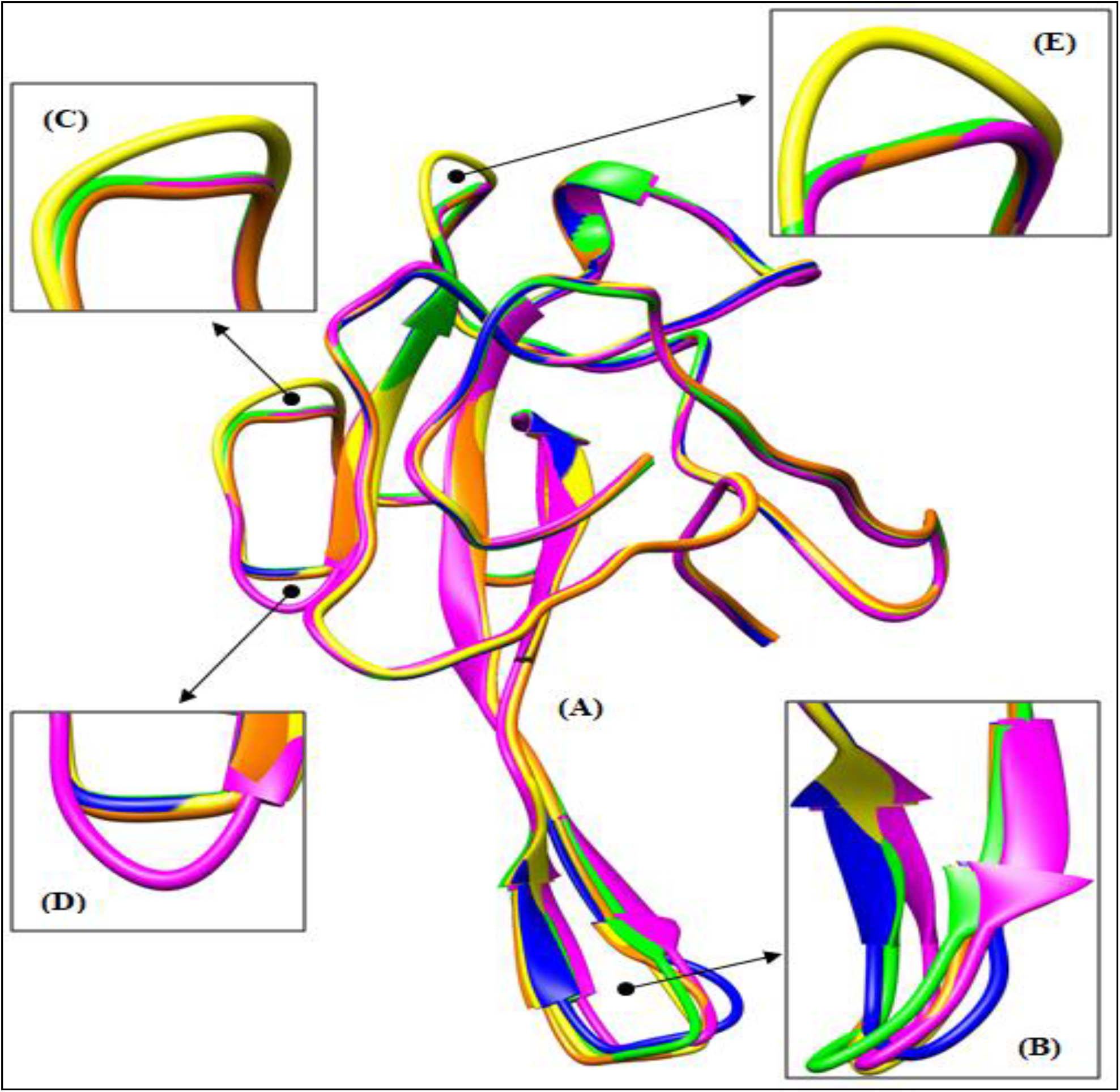
(A) Superimposition of CoV NTDs: SARS-CoV-2 (green), Pangolin-CoV MP789 (yellow), bat-SL-CoVZXC21 (orange), Pangolin-CoV GX-P5L (blue), and Bat-CoV RaTG13 (magenta). (B) Closer view of the distinct topological variations found within the β-hairpin region. (C) Flexible loop that was observed in Pangolin-CoV MP789. (D) Flexible loop that was observed in Bat-CoV RaTG13. (E) Flexible coil that was observed in Pangolin-CoV MP789.

Three other variations were observed. First, residues 123-125 in Pangolin-CoV MP789 showed a relaxed loop; however, it is not clear why this region is different as amino acids within this region are the same when compared to the CoV NTDs that do not present with this feature (Figure 6C; Figure S1). Second, residues 128-129 in Bat-CoV RaTG13 showed another relaxed loop. At residue position 128, the amino acid found was aspartic acid and is shared only with SARS-CoV-2 while the other CoV NTDs possess glutamic acid (Figure S1). Even though amino acid identity for this region is shared with SARS-CoV-2, only Bat-CoV RaTG13 displayed this relaxed loop (Figure 6D). Third, in the NTD for Pangolin-CoV MP789 residues 135-138 presented with a relaxed coil (Figure 6E). At residue 135, all other CoV NTDs have threonine while Pangolin-CoV MP789 possesses asparagine (Figure S1). Besides this difference, the remainder of this region holds the same amino acid identity for all NTDs.

Electrostatic potential maps revealed similar but innately visible differences in electrostatic surface patterns for each CoV NTD model (Figure S3). As expected, the positive charge was consolidated primarily throughout the β-hairpin region and upper binding groove. Interestingly, residue 80 within the 3/10-helices of the NTDs in both Pangolin-CoV GX-P5L and bat-SL-CoVZXC21 is lysine (Figure S1). Because of this extra positively charged amino acid, a small region of positive charge is visible on the top portion of their NTDs (Figure S3). This was not observed in other models; however, other NTDs that were not built presented with lysine at this site (Figure S1). The likely function and possible advantage of this residue was not determined in this study, but an investigation looking further into this residue is warranted.

## Discussion

Based on nucleotide and amino acid identity, the N gene and protein in SARS-CoV-2 share highest identity with the N gene and protein in Bat-CoV RaTG13; although, the N gene and protein found in Pangolin-CoV MP789 share nearly identical values. Each phylogenetic tree has Pangolin-CoV MP789 diverging prior to the SARS-CoV-2 and Bat-CoV RaTG13 sister group. This does not imply SARS-CoV-2 rose directly from either Pangolin-CoV MP789 or Bat-CoV RaTG13, but this is evidence of a close phylogenetic relationship. Additionally, it is unknown whether or not the common ancestors which gave rise to these lineages of CoVs are still in existence. At most, these data suggest a naturally intermingled bat and pangolin origin for SARS-CoV-2; however, the scope of this relationship is largely unclear. Both the N genes and proteins found among pangolin CoVs isolated between 2017-2018, bat-SL-CoVZXC21, and bat-SL-CoVZC45 expressed lower identities suggesting a slightly weaker relation to SARS-CoV-2. Perhaps these groups of viruses served as precursors that contributed to the evolutionary development of the SARS-CoV-2, Bat-CoV RaTG13, and Pangolin-CoV MP789 lineages. Alternatively, there is possibly an unsampled and circulating CoV in nature that is a missing link in the evolutionary history for SARS-CoV-2. Furthermore, the NTD and CTD in the SARS-CoV-2 N protein share highest identity with Bat-CoV RaTG13, followed by close identity with Pangolin-CoV MP789 as well. Slightly higher amino acid conservation than nucleotide conservation was observed for the entire N protein and both domains. Even though shared nucleotide identities were not extremely lower, this does suggest enough time has passed for silent mutations and/or genetic drift to begin taking minor effects on the N gene. In addition, the obvious redundancy in the genetic code likely contributes to this feature as well. Nucleotide data suggests the NTD is more conserved within the N gene. The NTD might be facing different selective pressures that are working to maintain less variability in nucleotide identity.

At the start of the N protein in SARS-CoV-2, Pangolin-CoV MP789, Bat-CoV RaTG13, bat-SL-CoVZXC21, and bat-SL-CoVZC45 all present with a double amino acid insertion. Though naturally occurring mutations are still possible, a recombination event could have occurred in these CoVs. Even so, sequences used in this study provide little evidence as to how this would have occurred. As previously suggested, perhaps an unsampled and widely circulating CoV in nature is responsible for introducing this type of insertion. Nonetheless, this feature reinforces a shared evolutionary history among these CoVs. In respect to the entire N protein, the NTR is disordered and functions in RNA-binding.^26^ The advantage of this insertion within the NTR is not clear, but it might help to further promote effective RNA-binding. For example, the insertions found within SARS-CoV-2, Pangolin-CoV MP789, Bat-CoV RaTG13, and bat-SL-CoVZXC21 possess arginine which is a positively charged amino acid that likely interacts with RNA. If so, this could have greater implications on the pathogenicity of CoVs possessing this insertion, such as in SARS-CoV-2. Experimental analysis could be an essential tool in determining the relative importance of this double insertion.

As for the amino acid residues specific to SARS-CoV-2, a prior study identified A267 as a positively selected site on an exposed loop within the CTD and is thought to evolve in a manner that might establish, maintain, or avoid binding of different host molecules.^13^ However, it appears little has been discussed about S37 which is found in the N-terminal region. According to the protein alignment, this residue is flanked by arginine and lysine residues. If a purpose for this mutation exists, it most likely associates with RNA binding. This serine could be phosphorylated, yielding a conformational change that allows flanking electropositive amino acids to bind with an RNA molecule. Although majority of phosphorylation sites are within the LKR region, a prior study investigating the SARS-CoV N protein found additional sites elsewhere, including the N-terminal region.^26,53^ Therefore, it is possible S37 could undergo phosphorylation to promote effective and high affinity RNA-binding. In support of this function, the prediction method that detected for residues likely to undergo phosphorylation indicated that S37 (score: 0.766) has a high potential of being phosphorylated. Experimental analysis is absolutely necessary to confirm this hypothesis. Alternatively, S37 lacks any functional role and is just the result of a random non-deleterious mutation. Lastly, both residues (S37 and A267) in SARS-CoV-2 were originally found by Zhang *et al*. 2020; therefore, sequence data in this study corroborates their findings.

Recombination analysis resulted in minimal evidence that recombination has occurred in the N gene. However, the N genes in Pangolin-CoV MP789 and Bat-CoV RaTG13 might be composed of a small fragment originally found in bat-SL-CoVZXC21. Considering that bat-SL-CoVZXC21 and Bat-CoV RaTG13 circulate among *Rhinolophus affinis*, perhaps a recombination event originated in that species and then made way into *Manis javanica*, resulting in a second recombination event between bat-SL-CoVZXC21 and Pangolin-CoV MP789. With bat-SL-CoVZXC21 serving as the source, this path of recombination is possible given the N gene is more conserved among Pangolin-CoV MP789 and Bat-CoV RaTG13. A recent study of the S gene in Pangolin-CoV MP789 suggested this specific gene was constructed by fragments found in either bat-SL-CoVZXC21 or bat-SL-CoVZC45, and from Bat-CoV RaTG13.^8^ Therefore, evidence of these CoVs having undergone recombination at some point is apparent and that nucleotides 485–565 for the N gene in SARS-CoV-2 was introduced from potential recombination events among Pangolin-CoV MP789, Bat-CoV RaTG13, and bat-SL-CoVZXC21.

Structurally, the NTD in Pangolin-CoV MP789 and Bat-CoV RaTG13 are similar to the NTD in SARS-CoV-2. This structural data reinforces the sequence-based phylogeny that reveals a shared evolutionary history for these CoVs. Among these NTDs, the region which displays the greatest topological variation is the β-hairpin motif. A recent study performed by Kang *et al.* 2020 elucidated the crystal structure for the NTD in SARS-CoV-2, and researchers discovered that when compared to the NTDs found in SARS-CoV, MERS-CoV and HCoV-OC43, the β-hairpin presented structural and topological variations.^54^ From this, it is understood these differences likely result in adaptive features that determine how specific CoV N proteins will bind to their own RNA genomes. Varying electrostatic surface potentials help support this concept. With varying distributions of positive charge, the mode of binding for the NTD will vary among N proteins. These differences in binding were also noted by Kang *et al.* 2020 in which they found the NTD in SARS-CoV-2 employs a unique binding pattern for ribonucleotides when compared to binding patterns for domains in SARS-CoV, MERS-CoV, and HCoV-OC43.^54^ For the NTDs found in SARS-CoV-2, Pangolin-CoV MP789, and Bat-CoV RaTG13 a structural relationship exists. However, electrostatic surfaces suggest unique modes of RNA-binding. Given that structures for the NTDs in Bat-CoV RaTG13 and Pangolin-CoV MP789 share greater similarity to the NTD in SARS-CoV-2 than with each other, it is likely these CoVs influenced the evolution leading up to the structural features observed in the SARS-CoV-2 NTD and how it binds to the genome. Also, an extended β-hairpin in SARS-CoV-2 might allow for a larger binding pocket that can accommodate RNA possessing higher ordered structure. Moreover, bending closer towards the binding site could permit a potentially tighter interaction with the genome. Discussed by Kang *et al*. 2020, they describe the N-terminal tail in the NTD as flexible and extending outward; therefore, this feature and the β-hairpin region might function together in allowing for a binding site that contains RNA of higher ordered structure.^54^ This could have further implications on how the genome is packaged when forming a helical nucleocapsid to make a complete virion. In addition, the topological variations observed in the β-hairpin region are likely not the result of intrinsic disorder. According to the disorder plot, this region presents with values that are slightly above the set threshold; however, these values are minimal enough that it is more likely residues comprising the β-hairpin region are ordered but perhaps moderately flexible in terms of accomplishing the most effective means of RNA-binding. Altogether, variations in β-hairpin topology could reflect how an N protein’s NTD interacts with a specified RNA genome.

Another recent study that identified epitopes in the SARS-CoV-2 N protein found the N protein in pangolin CoV failed to align with two of those epitope sequences.^55^ From this, the authors claim separations exist between functional adaptation and molecular evolution. There are variations between NTDs in SARS-CoV-2 and Pangolin-CoV MP789; despite their N proteins having a close evolutionary history. These differences in the NTD are likely reflective of adaptive functions in each N protein; therefore, it is likely there are features inherent only to the SARS-CoV-2 N protein that require further investigation. An example of such a specific feature would be the attributes associated with the β-hairpin motif. Given the N protein is immunogenic, knowing the sequence and structural differences can be useful for developing antigens in serological detection, antiviral medications, and vaccine candidates.^11,55^

In regard to the flexible loops and coil in Pangolin-CoV MP789 and Bat-CoV RaTG13, these features might also be inherent to their respective NTDs. Based on the data used to verify homology models, it seems these features are not errors as each model presented with ideal values for energy and stereochemical quality. For example, only asparagine is found at residue 135 in Pangolin-CoV MP789; therefore, this could affect spatial arrangements of subsequent amino acids and cause a looser coil to form (Figure S1). Though data is not shown, attempts at correcting these regions were done but resulted in lower stereochemical quality. In order to preserve a valid structure these flexible coils were not tampered with and kept as they were. Experimentally derived crystal structures for the NTD in these CoVs could help corroborate these findings.

It is clear CoVs originate in bats and can spillover into the human population through an intermediate host, such as with palm civets transmitting SARS-CoV.^13,14^ Unfortunately, this study does not provide evidence that pangolins are intermediate hosts for SARS-CoV-2 nor provide many insights into the chain of events leading up to the human spillover event. A major setback of this study is the lack of sequence data. With few sequences available from CoVs that are closely related to SARS-CoV-2, understanding the viruses’ origin is more difficult. However, by using the N gene and protein, this study was able to reinforce the existence of a close phylogenetic relationship among SARS-CoV-2, Bat-CoV RaTG13, and Pangolin-CoV MP789. Although the evolutionary history is similar, the N proteins in these CoVs seem to possess structural features that reflect upon adaptive functions specific to the CoVs they associate with. By continuing the surveillance of CoVs in bats and pangolins, it can help collect additional sequence information that can make greater sense of the relationship occurring between these CoVs. Ultimately, this would allow us to better understand the history of SARS-CoV-2 and what it could imply for the future should another novel CoV emerge causing a disease more dangerous than COVID-19.

## Acknowledgments

I want to thank all the previous scientists who have spent countless amounts of time researching SARS-CoV-2, along with their dedication to both understanding and fighting this ongoing pandemic. I also give thanks to Nataly Basora, Sabrina Krause, and Aziza Shemet-Pitcher for providing their insights on this project, along with helpful feedback during the initial construction of the manuscript. Furthermore, I especially want to thank Dr. Sharon Crary for providing a more technical review of the manuscript.

## References

1 He F, Deng Y, and Li W. (2020) “Coronavirus disease 2019: what we know?” Journal of Medical Virology. 2020. 92. Pg 719–725.

2 Brüssow H. (2020) “The Novel Coronavirus – A Snapshot of Current Knowledge.” Microbial Biotechnology. 13.3. Pg 607–612.

3 WHO: World Health Organization. (2020) “Coronavirus disease 2019 (COVID-19) Situation Report – 191.” Coronavirus disease (COVID-19) situation reports. World Health Organization.

4 Chen Y, Liu Q, and Guo D. (2020) “Emerging coronaviruses: Genome structure, replication, and pathogenesis.” Journal of Medical Virology. 92. Pg 418–423.

5 Gorbalenya AE, Baker SC, Baric RS, et al. (2020) “The species *Severe acute respiratory syndrome-related coronavirus*: classifying 2019-nCoV and naming it SARS-CoV-2.” Nature Microbiology. 5. Pg 536–544.

6 Lu R, Zhao X, Li J, et al. (2020) “Genomic characterisation and epidemiology of 2019 novel coronavirus: implications for virus origins and receptor binding.” The Lancet. 395. Pg 565–574.

7 Wu A, Peng Y, Huang B, et al. (2020) “Genome Composition and Divergence of the Novel Coronavirus (2019-nCoV) Originating in China.” Cell Host & Microbe. 27.3. Pg 325–328.

8 Liu P, Jiang J-Z, Wan X-F, et al. (2020) “Are pangolins the intermediate host of the 2019 novel coronavirus (SARS-CoV-2)?” PLoS Pathogens. 16.5. Pg 1–13.

9 Wong MC, Sara JC, Ajami NJ, et al. (2020) “Evidence of recombination in coronaviruses implicating pangolin origins of nCoV-2019.” bioRxiv. Preprint.

10 Andersen KG, Rambaut A, Lipkin WI, et al. (2020) “The proximal origin of SARS-CoV-2.” Nature Medicine. 26. Pg 450–452.

11 Zhang T, Wu Q, and Zhang Z. (2020) “Probable Pangolin Origin of SARS-CoV-2 Associated with the COVID-19 Outbreak.” Current Biology. 30. Pg 1346–1351.

12 Tang X, Wu C, Li X, et al. (2020) “On the origin and continuing evolution of SARS-CoV-2.” National Science Review. 7.6. Pg 1012–1023.

13 Cagliani R, Forni D, Clerici M, et al. (2020) “Computational inference of selection underlying the evolution of the novel coronavirus, SARS-CoV-2.” Journal of Virology. 94. Pg. 411–420.

14 Vijaykrishna D, Smith GJ, Zhang JX, et al. (2007) “Evolutionary insights into the ecology of coronaviruses.” Journal of Virology. 81.8. Pg 4012–4020.

15 Cui J, Li F, and Shi Z. (2019) “Origin and evolution of pathogenic coronaviruses.” Nature Reviews Microbiology. 17. Pg 181–192.

16 Ji W, Wang W, Zhao X, et al. (2020) “Cross‐species transmission of the newly identified coronavirus 2019‐nCoV.” Journal of Medical Virology. 92. Pg 433–440.

17 Liu Z, Xiao X, Wei X, et al. (2020). “Composition and divergence of coronavirus spike proteins and host ACE2 receptors predict potential intermediate hosts of SARS-CoV-2.” Journal of Medical Virology. 92. Pg 595–601.

18 Dabravolski SA and Kavalionak YK. (2020) “SARS‐CoV‐2: Structural diversity, phylogeny, and potential animal host identification of spike glycoprotein.” Journal of Medical Virology. Pg 1–5.

19 Zheng J. (2020). “SARS-CoV-2: an Emerging Coronavirus that Causes a Global Threat.” International Journal of Biological Sciences. 16.10. Pg 1678–1685.

20 Tok TT and Tatar G. (2017) “Structures and Functions of Coronavirus Proteins: Molecular Modeling of Viral Nucleoprotein.” International Journal of Virology & Infectious Disease. 2.1. Pg 1–7.

21 Gui M, Liu X, Guo D, et al. (2017) “Electron microscopy studies of the coronavirus ribonucleoprotein complex.” Protein & Cell. 8.3. Pg 219–224.

22 Lo CY, Tsai TL, Lin CN, et al. (2019) “Interaction of coronavirus nucleocapsid protein with the 5’- and 3’-ends of the coronavirus genome is involved in genome circularization and negative-strand RNA synthesis.” The FEBS Journal. 286.16. Pg 3222–3239.

23 Neuman BW and Buchmeier MJ. (2016) “Supramolecular Architecture of the Coronavirus Particle.” Advances in Virus Research. 96. Pg 8–9.

24 Ahmed SF, Quadeer AA, and McKay MR. (2020) “Preliminary Identification of Potential Vaccine Targets for the COVID-19 Coronavirus (SARS-CoV-2) Based on SARS-CoV Immunological Studies.” Viruses. 12(3).254. Pg 1–15.

25 Liu SJ, Leng CH, Lien SP, et al. (2006) “Immunological characterizations of the nucleocapsid protein based SARS vaccine candidates.” Vaccine. 24.16. Pg 3100–3108.

26 McBride R, van Zyl M, and Fielding BC. (2014) “The coronavirus nucleocapsid is a multifunctional protein.” Viruses. 6.8. Pg 2991–3018.

27 Chang CK, Hsu YL, Chang YH, et al. (2009) “Multiple Nucleic Acid Binding Sites and Intrinsic Disorder of Severe Acute Respiratory Syndrome Coronavirus Nucleocapsid Protein: Implications for Ribonucleocapsid Protein Packaging.” Journal of Virology. 85.5. Pg 2255–2264.

28 Saikatendu KS, Joseph JS, Subramanian V, et al. (2007) “Ribonucleocapsid formation of severe acute respiratory syndrome coronavirus through molecular action of the N-terminal domain of N protein.” Journal of Virology. 81.8. Pg 3913–3921.

29 Huang Q, Yu L, Petros AM, et al. (2004) “Structure of the N-terminal RNA-binding domain of the SARS CoV nucleocapsid protein.” Biochemistry. 43.20. Pg 6059–6063.

30 Dilucca M, Forcelloni S, Georgakilas AG, et al. (2020) “Temporal evolution and adaptation of SARS-COV 2 codon usage.” bioRxiv. Preprint.

31 Yu IM, Oldham ML, Zhang J, et al. (2006) “Crystal Structure of the Severe Acute Respiratory Syndrome (SARS) Coronavirus Nucleocapsid Protein Dimerization Domain Reveals Evolutionary Linkage between Corona- and Arteriviridae.” The Journal of Biological Chemistry. 281.25. Pg 17134–17139.

32 Campanella JJ, Bitincka L, and Smalley J. (2003) “MatGAT: An application that generates similarity/identity matrices using protein or DNA sequences.” BMC Bioinformatics. 4.29. Pg 1–4.

33 Edgar RC. (2004) “MUSCLE: multiple sequence alignment with high accuracy and high throughput.” Nucleic Acids Research. 32.5. Pg. 1792–1797.

34 Kumar S, Stecher G, Li M, et al. (2018) “MEGA X: Molecular Evolutionary Genetics Analysis across computing platforms.” Molecular Biology and Evolution 35. Pg 1547–1549.

35 Whelan S and Goldman N. (2001) “A general empirical model of protein evolution derived from multiple protein families using a maximum-likelihood approach.” Molecular Biology and Evolution. 18. Pg 691–699.

36 Lole KS, Bollinger RC, Paranjape RS, et al. (1999) “Full-length human immunodeficiency virus type 1 genomes from subtype C-infected seroconverters in India, with evidence of intersubtype recombination.” Journal of Virology. 73.1. Pg 152–160.

37 Ishida T and Kinoshita K. (2007). “PrDOS: prediction of disordered protein regions from amino acid sequence.” Nucleic Acids Research. 35. Pg 460–464.

38 Erdős G and Dosztányi Z. (2020) “Analyzing Protein Disorder with IUPred2A.” Current Protocols in Bioinformatics. 70.1. Pg 1–15.

39 Mészáros B, Erdős G, and Dosztányi Z. (2018) “IUPred2A: context-dependent prediction of protein disorder as a function of redox state and protein binding.” Nucleic Acids Research. 46.1. Pg 329–337.

40 Jones DT and Cozzetto D. (2015) “DISOPRED3: precise disordered region predictions with annotated protein-binding activity.” Bioinformatics. 31.6. Pg 857–863.

41 Cleveland SB, Davies J, and McClure MA. (2011) “A Bioinformatics Approach to the Structure, Function, and Evolution of the Nucleoprotein of the Order Mononegavirales.” PLoS ONE. 6.5. Pg 1–13.

42 Fiser A and Šali A. (2003) “MODELLER: generation and refinement of homology-based protein structure models.” Methods in Enzymology. 374. Pg 463–493.

43 Eswar N, Webb B, Marti-Renom MA, et al. (2006) “Comparative protein structure modeling using Modeller.” Current Protocols in Bioinformatics. 5. Pg 1–47.

44 Bowie JU, Lüthy R, and Eisenberg D. (1991) “A method to identify protein sequences that fold into a known three-dimensional structure.” Science. 253.5016. Pg 164–170.

45 Lüthy R, Bowie JU, and Eisenberg D. (1992) “Assessment of protein models with three-dimensional profiles.” Nature. 356.6364. Pg 83–85.

46 Colovos C and Yeates TO. (1993) “Verification of protein structures: patterns of nonbonded atomic interactions.” Protein Science. 2.9. Pg 1511–1519.

47 Laskowski RA, Rullmann JAC, MacArthur MW, et al. (1996) “AQUA and PROCHECK-NMR: Programs for checking the quality of protein structures solved by NMR.” Journal of Bimolecular NMR. 8. Pg 477–486.

48 Laskowski RA, MacArthur MW, Moss DS, et al. (1993) “PROCHECK: a program to check the stereochemical quality of protein structures.” Journal of Applied Crystallography. 26. Pg 283–291.

49 Pettersen EF, Goddard TD, Huang CC, et al. (2004) “UCSF Chimera-a visualization system for exploratory research and analysis.” Journal of Computational Chemistry. 25.13. Pg 1605–1612.

50 Meng EC, Pettersen EF, Couch GS, et al. (2006) “Tools for integrated sequence-structure analysis with UCSF Chimera.” BMC Bioinformatics. 7.339. Pg 1–10.

51 Johnson MS, Sutcliffe MJ, and Blundell TL. (1990) “Molecular anatomy: phyletic relationships derived from three-dimensional structures of proteins.” Journal of Molecular Evolution. 30.1. Pg 43–59.

52 Krissinel E and Henrick K. (2004) “Secondary-structure matching (SSM), a new tool for fast protein structure alignment in three dimensions.” Acta Crystallographica Section D. 60.12(1). Pg 2256–2268.

53 Lin L, Shao J, Sun M, et al. (2007) “Identification of phosphorylation sites in the nucleocapsid protein (N protein) of SARS-coronavirus.” International Journal of Mass Spectrometry. 268.2. Pg 296–303.

54 Kang S, Yang M, Hong Z, et al. (2020) “Crystal structure of SARS-CoV-2 nucleocapsid protein RNA binding domain reveals potential unique drug targeting sites.” Acta Pharmaceutica Sinica B. [published online ahead of print, 2020 Apr 20].

55 Tilocca B, Soggiu A, Sanguinetti M, et al. (2020) “Comparative computational analysis of SARS-CoV-2 nucleocapsid protein epitopes in taxonomically related coronaviruses.” Microbes and Infection. 22.4-5. Pg 188–194.

