## Supplemental Figure 1 for "Using nucleocapsid proteins to investigate the relationship between SARS-CoV-2 and closely related bat and pangolin coronaviruses"

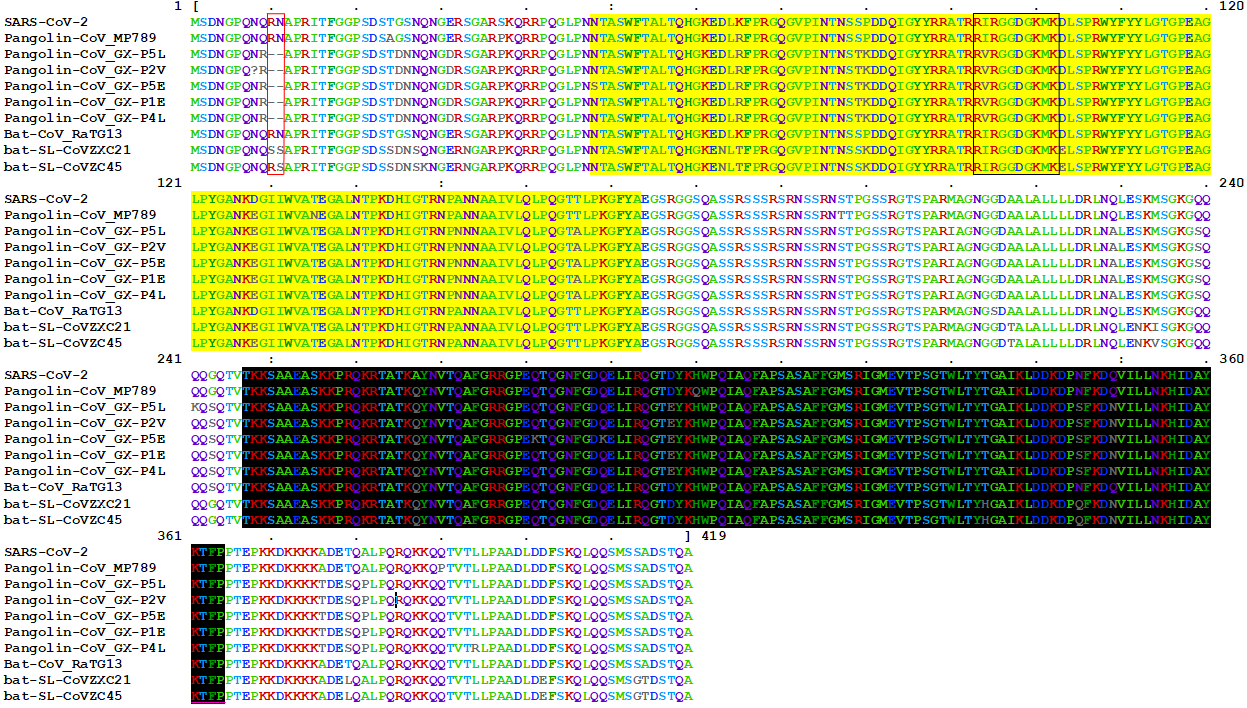
Figure S1. Multiple sequence alignment generated at the amino acid level for the N protein. The alignment was generated with MUSCLE on MEGE-X and visualized using MView on the EMBL-EBI online server: <https://www.ebi.ac.uk/Tools/msa/mview/>. Colors represent amino acids of similar identity whereas grey amino acids indicate a difference found in the alignment. The red box indicates the double amino acid insertion region and the black box indicates the amino acids present within the β-hairpin for each N protein. The region highlighted in yellow covers the NTD and the region highlighted in black covers the CTD.
