## Supplemental Figure 2 for "Using nucleocapsid proteins to investigate the relationship between SARS-CoV-2 and closely related bat and pangolin coronaviruses"

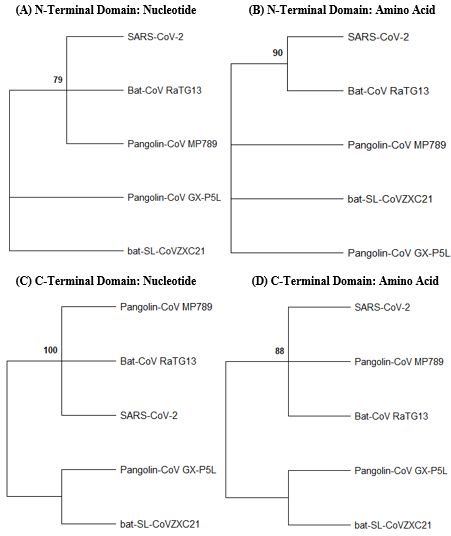
 Figure S2. Phylogenetic analysis of NTD and CTD sequences (nucleotide and amino acid) depicting evolutionary relationships observed among SARS-CoV-2, pangolin CoVs, and closely related	bat CoVs. (A) Phylogeny was inferred using the Jukes-Cantor substitution model. A log likelihood	of -794.07 was determined with bootstrap values calculated out of 500 replicates and nodes	with < 70.00% support collapsed. (B) The phylogeny was inferred using the JTT matrix-based substitution model. A log likelihood of -440.56 was determined with bootstrap values calculated out of 500 replicates and nodes with < 70.00% support collapsed. (C) Phylogeny was inferred using	the Jukes-Cantor substitution model. A log likelihood of -870.56 was determined with bootstrap values calculated out of 500 replicates and nodes with < 70.00% support collapsed. (D) The phylogeny was inferred using the JTT matrix-based substitution model. A log likelihood of -400.46 was determined with bootstrap values calculated out of 500 replicates and nodes with < 70.00% support collapsed.
