## Supplemental Figure 3 for "Using nucleocapsid proteins to investigate the relationship between SARS-CoV-2 and closely related bat and pangolin coronaviruses"

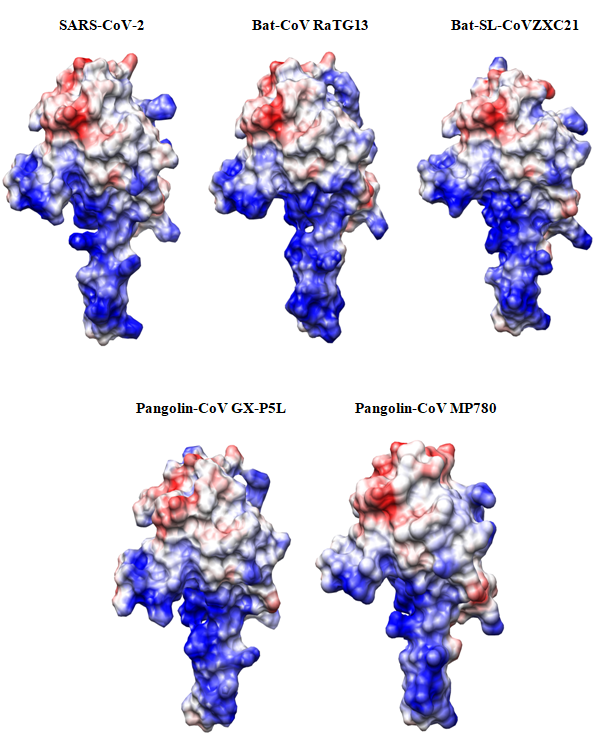
Figure S3. Electrostatic surface potential maps of the NTD in SARS-CoV-2, Bat-CoV RaTG13, bat-SL-CoVZXC21, Pangolin-CoV GX-5PL, and Pangolin-CoV MP789. Blue indicates regions of positive charge and red indicates regions of negative charge. White indicates regions of neutral charge.
