## Supplemental Table 1 for "Using nucleocapsid proteins to investigate the relationship between SARS-CoV-2 and closely related bat and pangolin coronaviruses"

| **CoV Species** | **Nucleotide Reference** | **Protein Reference** | **Host** | **Year of Isolation** | **Geographic Location** |
| --- | --- | --- | --- | --- | --- |
| SARS-CoV-2 | NC_045512.2 | YP_009724397.2 | *Homo sapiens* | 2019 | Hubei |
| Pangolin-CoV MP789 | MT121216.1 | QIG55953.1 | *Manis javanica* | 2019 | Guangdong |
| Pangolin-CoV GX-P5L | MT040335.1 | QIA48639.1 | *Manis javanica* | 2017 | Guangxi |
| Pangolin-CoV GX-P2V | MT072864.1 | QIQ54056.1 | *Manis javanica* | 2018 | Guangxi |
| Pangolin-CoV GX-P5E | MT040336.1 | QIA48648.1 | *Manis javanica* | 2017 | Guangxi |
| Pangolin-CoV GX-P1E | MT040334.1 | QIA48630.1 | *Manis javanica* | 2017 | Guangxi |
| Pangolin-CoV GX-P4L | MT040333.1 | QIA48621.1 | *Manis javanica* | 2017 | Guangxi |
| Bat-CoV RaTG13 | MN996532.1 | QHR63308.1 | *Rhinolophus affinis* | 2013 | Yunnan |
| bat-SL-CoVZXC21 | MG772934.1 | AVP78049.1 | *Rhinolophus affinis* | 2015 | Zhejiang |
| bat-SL-CoVZC45 | MG772933.1 | AVP78038.1 | *Rhinolophus sinicus* | 2017 | Zhejiang |

Table S1. The CoVs and their corresponding nucleocapsid gene and protein sequences used in this study, along with the host, year of isolation, and geographic location (i.e. all Chinese provinces or autonomous regions).
