## Supplemental Table 2 for "Using nucleocapsid proteins to investigate the relationship between SARS-CoV-2 and closely related bat and pangolin coronaviruses"

| **CoV Species** | **Nucleotide Identity (%)** | **Amino Acid Identity (%)** |
| --- | --- | --- |
| Pangolin-CoV MP789 | 96.2 | 97.9 |
| Pangolin-CoV GX-P5L | 91.0 | 93.8 |
| Pangolin-CoV GX-P2V | 91.0 | 93.8 |
| Pangolin-CoV GX-P5E | 90.8 | 93.3 |
| Pangolin-CoV GX-P1E | 91.0 | 94.0 |
| Pangolin-CoV GX-P4L | 91.0 | 93.3 |
| Bat-CoV RaTG13 | 96.9 | 99.0 |
| bat-SL-CoVZXC21 | 91.2 | 94.3 |
| bat-SL-CoVZC45 | 91.1 | 94.3 |

Table S2. Nucleocapsid gene and protein identities compared against SARS-CoV-2.
