## Supplemental Table 5 for "Using nucleocapsid proteins to investigate the relationship between SARS-CoV-2 and closely related bat and pangolin coronaviruses"

| **CoV Species** | **DOPE Score** | **Verify-3D** | **ERRAT** | **PROCHECK** |
| --- | --- | --- | --- | --- |
| SARS-CoV-2 | -11,146.10547 | 96.83% | 90.678 | 95.90% |
| Bat CoV RaTG13 | -11,123.69824 | 90.40% | 91.5254 | 96.90% |
| bat-SL-CoVZXC21 | -11,250.74902 | 94.44% | 91.5254 | 97.00% |
| Pangolin CoV GX-P5L | -11,269.37793 | 100.00% | 90.678 | 97.00% |
| Pangolin CoV MP789 | -11,143.73145 | 97.62% | 93.2203 | 93.90% |

Table S3. Final energy and stereochemical values for each NTD homology model.
